# CDK12 is Necessary to Promote Epidermal Differentiation through Transcription Elongation

**DOI:** 10.1101/2021.06.01.446524

**Authors:** Jingting Li, Manisha Tiwari, Yifang Chen, George L. Sen

## Abstract

Proper differentiation of the epidermis is essential to prevent water loss and to protect the body from the outside environment. Perturbations in this process can lead to a variety of skin diseases that impacts 1 in 5 people. While transcription factors that control epidermal differentiation have been well characterized, other aspects of transcription control such as elongation are poorly understood. Here we show that of the two cyclin dependent kinases (CDK12 and CDK13), that are known to regulate transcription elongation, only CDK12 is necessary for epidermal differentiation. Depletion of CDK12 led to loss of differentiation gene expression and absence of skin barrier formation in regenerated human epidermis. CDK12 binds to genes that code for differentiation promoting transcription factors (*GRHL3, KLF4*, and *OVOL1*) and is necessary for their elongation. CDK12 is necessary for elongation by promoting Ser2 phosphorylation on the C-terminal domain of RNA polymerase II and the binding of the elongation factor SPT6 to target genes. Our results suggest that control of transcription elongation by CDK12 plays a prominent role in adult cell fate decisions.

## INTRODUCTION

Proper regulation of gene expression is not only essential for the development of an organism but also for cells to respond to stress, environmental stimulus as well as balance growth and differentiation in the adult state. Dysregulation of gene expression has long been known to contribute to human disease most notably in tumorigenesis^1,2^. Studies on regulation of gene expression have been primarily focused on transcription factors but they represent only one aspect of transcription. Transcription driven by RNA polymerase II (Pol II) is regulated at multiple levels including initiation, pausing, elongation, and termination with each of these stages regulated by specific cyclin dependent kinases (CDKs) and their respective cyclins^3^. The preinitiation complex (PIC) is formed when general transcription factors (GTFs) are assembled at promoters and cause the recruitment of Pol II^4^. Once recruited, CDK7/Cyclin H, which is associated with TFIIH (GTF), phosphorylates the C-terminal domain of Pol II on serine 5 (ser5) and serine 7 (ser7) to promote initiation^5-7^. Ser5 phosphorylation allows Pol II release from the PIC and transcription of short nascent RNA which allows binding of the DRB sensitivity-inducing factor (DSIF: SPT4 and SPT5) and negative elongation factor (NELF)^8-10^. Binding of DSIF/NELF leads to Pol II pausing typically after transcribing 20-60bps of RNA^11-13^. Release of paused Pol II into productive transcription occurs when positive elongation factor b (P-TEFb: CDK9 and Cyclin T) phosphorylates DSIF, NELF, and serine 2 (Ser2) of Pol II’s CTD^14^. Phosphorylation of SPT5 and NELF results in the dissociation of NELF from Pol II^15^. Without the inhibitory NELF, the PAF complex and SPT6 can associate with Pol II to facilitate transcription elongation^10,16^. SPT6 is a histone chaperone that interacts with both histones H3 and H4 to disassemble and reassemble nucleosomes to allow passage of Pol II during elongation^4,17^

While the role of CDK7/cyclin H and CDK9/cyclin T in Pol II initation and pause-release respectively has been well studied, the CDKs involved in promoting transcription elongation are much less characterized. Recent studies have shown that CDK12 and CDK13 perform the majority of Ser2 phosphorylation on Pol II’s CTD to regulate elongation^18,19^. It is currently unclear whether CDK12 or CDK13 have any roles in regulating cell fate decisions and tissue homeostasis. These questions have not been addressed due to the embryonic lethal phenotype in mice of knockout of these genes^20,21^. Understanding the roles of these CDKs in tissue development/homeostasis is especially pertinent as many of these are dysregulated in diseases such as cancer^22,23^.

The human epidermis is a fast turnover tissue which constantly relies on stem and progenitor cells residing in the basal layer to continuously supply differentiated cells to form the barrier of our skin^24^. Alterations in the differentiation process that leads to defective barrier formation (disrupted stratum corenum) can cause a variety of skin diseases which afflicts ∼20% of the population^25^. Proper keratinocyte differentiation requires an exit out of the cell cycle and induction of the differentiation gene expression program. Most studies on epidermal differentiation have focused on identifying and characterizing transcription or epigenetic factors necessary for this process. Work from our lab and others have shown that key factors such as ZNF750, KLF3, KLF4, MAF, MAFB, GRHL1, OVOL1, GRHL3, EHF, BRD4, JMJD3 and CEBP alpha/beta are essential for differentiation^26-36^. Intriguingly, our recent work demonstrated that ∼30% of epidermal differentiation genes already contained paused Pol II at its transcriptional start site (TSS) in stem and progenitor cells^37^. Many of these poised differentiation genes with paused Pol II binding coded for the differentiation promoting transcription factors described above. Upon induction of differentiation, Pol II is released into productive elongation allowing for the expression of these transcription factors which then turns on the rest of the epidermal differentiation gene expression program. A small RNAi screen of pause release/elongation factors identified SPT6 and PAF1 to be the key factors involved in the elongation of the differentiation specific transcription factors as well as structural differentiation genes^37^. Loss of function of SPT6 or PAF1 led to inhibition of epidermal differentiation and accumulation of Pol II at the TSS of differentiation genes. These results suggest that regulation of transcription elongation plays a prominent role in cell fate decisions and suggests that the CDKs that regulate this step may potentially be important for this process.

Here, we show that of the CDKs that regulate transcription elongation, only CDK12 loss blocked differentiation of regenerated human skin. CDK12 bound to over 8,000 genes primarily localized to their TSS. CDK12 also associated with over half of the genes that SPT6 bound.

These genes include key differentiation promoting transcription factors such as *OVOL1, GRHL3*, and *KLF4* as well as differentiation specific structural genes such as *TGM1*. CDK12 is necessary for Pol II Ser2 phosphorylation of its bound genes. In the absence of CDK12, Pol II Ser2 phosphorylation is lost on critical epidermal differentiation genes, which results in Pol II depletion from the gene body and buildup at the TSS. This loss of elongation causes decreased expression of crucial differentiation genes, which in turn leads to blockade of terminal epidermal differentiation. Without CDK12, elongation factors such as SPT6 are unable to bind. These results highlight the prominent role that transcription elongation factors have on somatic tissue differentiation.

### CDK12 but not CDK13 is Necessary to Promote Epidermal Differentiation

To test whether the CDKs that regulate transcription elongation (CDK12 and CDK13), have any impact on epidermal function, each was knocked down in primary human keratinocytes using siRNAs. The knockdown cells were placed in high calcium and full confluence for 3 days to induce differentiation. Interestingly, loss of CDK13 did not impact expression of epidermal differentiation genes (SFIG 1A). In contrast, CDK12 knockdown [(validated through 2 distinct siRNAs (CDK12-Ai and CDK12-Bi) targeting different regions of the gene)] blocked expression of critical differentiation structural genes such as *KRT10, FLG, LOR, ABCA12*, and *TGM1* (Figure 1A). Notably, mutations in genes such as *KRT10, FLG, ABCA12*, and *TGM1* lead to severe skin diseases such as epidermolytic hyperkeratosis, ichthyosis vulgaris, harlequin ichthyosis, and congenital ichthyosis respectively^38-41^.

**Figure 1.**
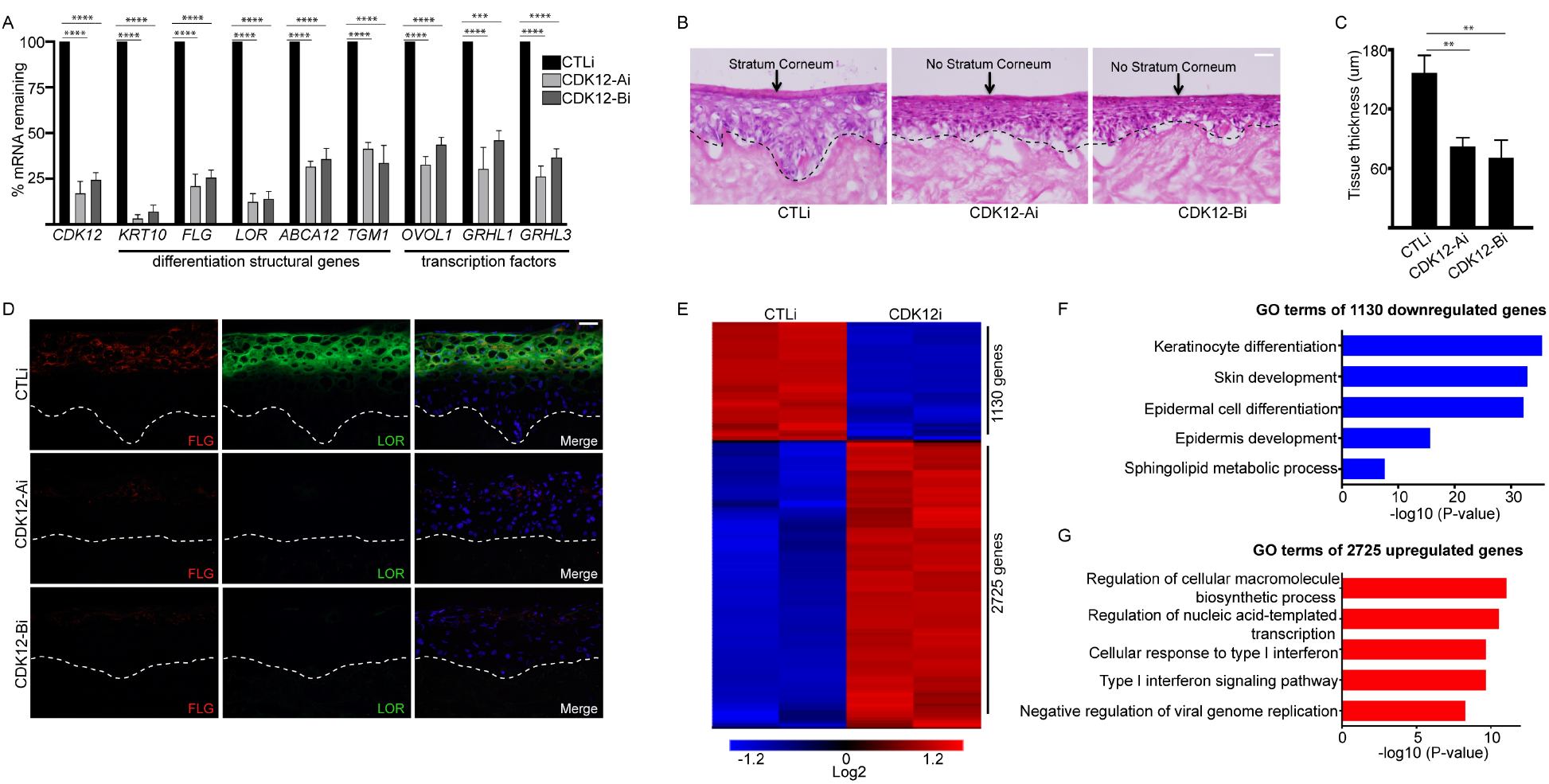
CDK12 is Necessary for Epidermal Differentiation. **(A)** Primary human keratinocytes were knocked down with control (CTLi) or CDK12 (CDK12-Ai and CDK12-Bi are distinct siRNAs that target 2 different regions of the gene) siRNAs and placed in differentiation conditions for 3 days (full confluence and 1.2mM calcium). The mRNA levels of each gene (X-axis) were measured using RT-qPCR. QPCR results were normalized to *L32* levels. **(B)** Regenerated human skin using three-dimensional organotypic cultures made from CTLi or CDK12i cells were harvested after 4 days of culture. Hematoxylin and eosin staining of regenerated CTLi and CDK12i human skin. The dashed black lines denote the basement membrane zone and scale bar=20μm. **(C)** Quantification of tissue thickness from (B). **(D)** Immunostaining of late differentiation proteins filaggrin (FLG: red) and loricrin (LOR: green) are shown (Day 4 regenerated human skin). Nuclei are shown in blue (Hoechst staining). The dashed white lines denote the basement membrane zone and scale bar=20μm. **(E)** RNA-Seq analysis of CTL and CDK12 knockdown primary human keratinocytes differentiated for 3 days (full confluence and 1.2mM calcium). RNA-Seq results were performed in duplicates. 2,725 genes were upregulated (red) and 1,130 genes were downregulated (blue) upon CDK12 knockdown. Heat map is shown in Log2 scale. **(F)** Gene ontology (GO) terms for the 1,130 genes downregulated in CDK12i cells. **(G)** GO terms for the 2,725 genes with increased expression upon CDK12 depletion. N=3 independent experiments for Figure 1 unless otherwise indicated. Mean values are shown with error bars=SD. ** ***p*** <0.01, \*\*\****p*** <0.001, \*\*\*\****p*** <0.0001 (t-test). Source data for Figure 1A, 1C can be found in Figure 1-Source Data 1.

Differentiation promoting transcription factors that we previously showed to be controlled by transcription elongation were also reduced in expression in CDK12i cells^37^ (Figure 1A). Control and knockdown primary human keratinocytes were also used to regenerate human skin by seeding the cells on devitalized human dermis^42-44^. This allows the cells to establish cell-cell and cell-basement membrane contact to allow proper growth, differentiation, and stratification in 3 dimensions. This technology has been used to cure patients of junctional epidermolysis bullosa through the autologous transplantation of transgenic regenerated human skin^45,46^. Similar to cells cultured in 2D, CDK13 knockdown did not alter skin proliferation, differentiation, or morphology (SFIG 1B-E). CDK12 loss blocked differentiation as evidenced by a lack of stratum corneum formation (Figure 1B). The terminally differentiated layer of the epidermis did not form due to an absence of filaggrin (FLG) and loricrin (LOR) protein expression (Figure 1D). Early differentiation protein (keratin 10: K10) expression was also downregulated (SFIG 2A). The proliferative capacity of the basal layer was also compromised as evidenced by the loss of ki67 positive cells which resulted in a hypoplastic tissue (Figure 1C, SFIG 2A-B). To understand the genes that CDK12 is regulating on a genome-wide level, RNA-sequencing (RNA-Seq) was performed on CTLi and CDK12i cells cultured in differentiation conditions for 3 days. 1,130 genes (p-value < 0.05 and ≥ 2-fold change) decreased in expression upon CDK12 depletion, which were enriched in genes involved in keratinocyte differentiation, skin development, and sphingolipid metabolic process (Figure 1E-F, Supplementary Table 1). 2,725 genes were upregulated upon CDK12 knockdown, which were enriched in regulation of cellular macromolecule biosynthetic process and cellular response to type I interferon (Figure 1E, 1G, Supplementary Table 1). The increased expression of interferon related genes may be a response to the barrier loss in CDK12i cells. A comparison with the differentiation gene expression signature (differentially expressed genes upon epidermal differentiation^30^) showed that 33.3% of the CDK12 gene signature overlapped (SFIG 2C). Importantly these 1,285 overlapped genes were enriched for gene ontology (GO) terms such as epidermis development suggesting that CDK12 is vital for epidermal differentiation and may control this process through elongation (SFIG 2D).

### CDK12 Binds and Regulates Epidermal Differentiation Genes

To determine whether CDK12 directly regulated the expression of epidermal differentiation genes, we performed ChIP-Seq on CDK12 in differentiated human keratinocytes. CDK12 bound to 12,230 peaks with the vast majority (88%) of the binding sites mapping back to genic regions (promoter, intron, exon, 5’UTR, TTS, and 3’UTR) (Figure 2A). The 12,230 CDK12 bound sites mapped back to 8,630 genes which were enriched for GO terms such as “negative regulation of transcription” and “ubiquitin dependent protein catabolic process” (Figure 2B, Supplementary Table 2). Since CDK12 has previously^47^ been implicated in regulating transcription elongation through Pol II, we compared our previously^37^ generated Pol II ChIP-Seq data to CDK12. The CDK12 and Pol II binding profiles were similar at CDK12 bound genes (Figure 2C-D). There were also notable differences in binding as CDK12 binding was primarily concentrated at the TSS of genes whereas Pol II binding also had another peak near the transcription end site (TES) (Figure 2D). 40.6% of the genes increased upon CDK12 loss were bound by CDK12 and were enriched in GO terms such as positive regulation of cell morphogenesis and negative regulation of epithelial cell proliferation (Figure 2E-F). 490 (43.4%) of the genes decreased upon CDK12 knockdown were also bound by CDK12 and were enriched for establishment of skin barrier and positive regulation of epidermis development GO terms (Figure 2E,2G). This included genes such that coded for key transcription factors (*GRHL3, OVOL1, KLF4*) that promotes epidermal differentiation as well as structural differentiation genes (*TGM1*) suggesting that CDK12 may regulate their elongation (Figure 2H, SFIG 3A-C).

**Figure 2.**
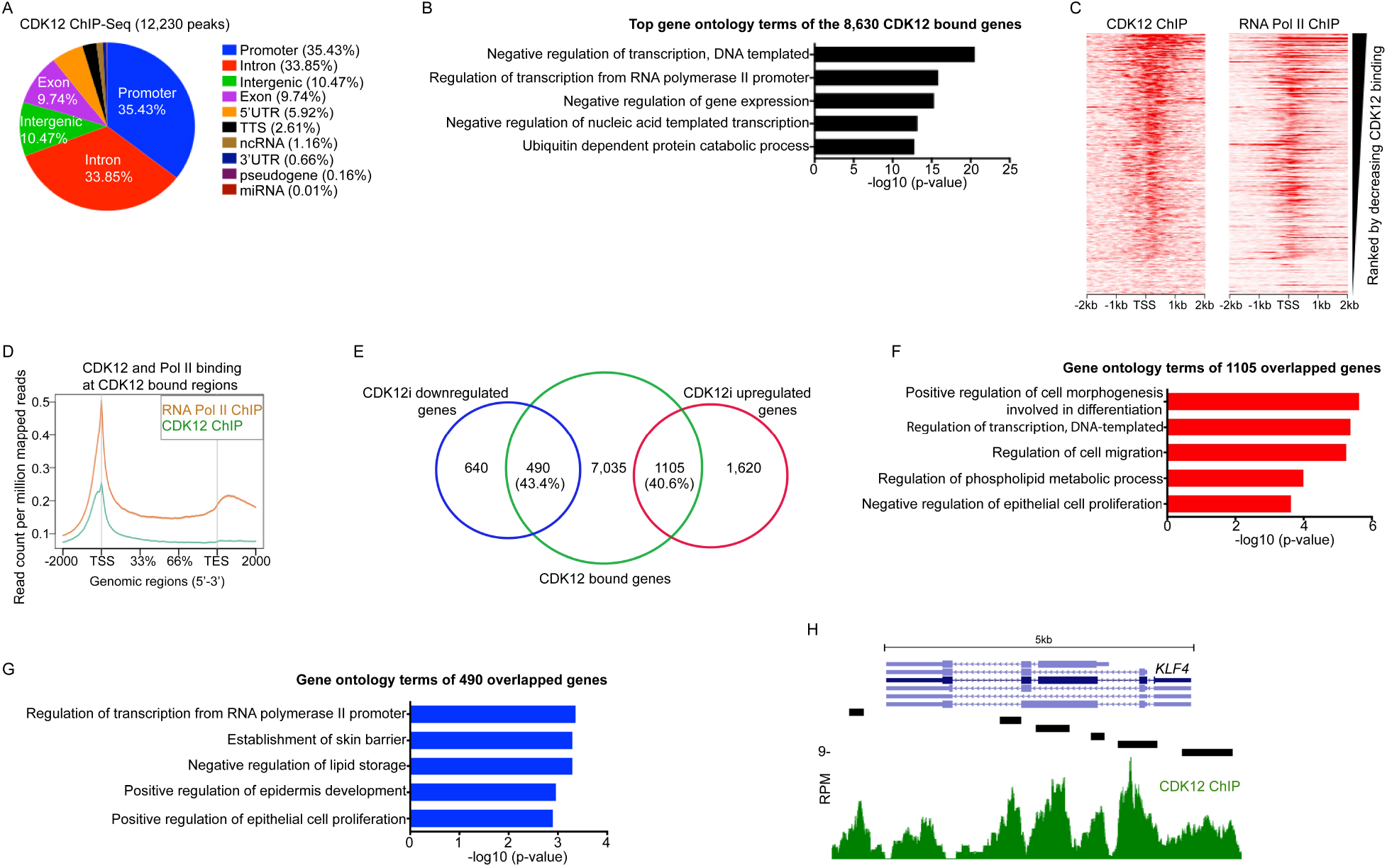
CDK12 Binds to Genes Coding for Epidermal Differentiation Proteins. **(A)** Genomic localization of the 12,230 CDK12 bound peaks. The percent of CDK12 binding to each genomic region is shown. CDK12 ChIP-Seq was performed in primary human keratinocytes cultured in differentiation conditions (full confluence and 1.2mM calcium: day 3). N=2. **(B)** Gene ontology terms of the 8,630 genes that CDK12 binds. **(C)** Heatmap of CDK12 and RNA Pol II ChIP-Seq (day 3 differentiated cells) on sites bound by CDK12. X-axis shows - 2kb to +2kb from the TSS. Heatmap is shown ranked by decreasing CDK12 binding. **(D)** Metagene plot of CDK12 (teal) and RNA Pol II (orange) ChIP-Seq reads at CDK12 bound regions. Y-axis is shown as read count per million reads and X-axis is distance along CDK12 bound genes. TSS is transcription start site and TES is transcription end site. **(E)** Venn diagram of CDK12 bound genes (CDK12 ChIP-Seq) with genes upregulated or downregulated upon CDK12 depletion. **(F)** GO terms of the CDK12 bound genes that overlap with genes increased upon CDK12 loss. **(G)** GO terms of the 490 decreased genes upon CDK12 knockdown that overlap with CDK12 bound genes. **(H)** Gene track of *KLF4*. CDK12 ChIP-Seq is shown in green. Y-axis shows reads per million (RPM) and black bar over gene track represents significant peaks. X-axis shows position along gene.

### CDK12 Promotes Elongation of Differentiation Genes through Pol II Ser2 Phosphorylation and Binding of the Elongation Factor SPT6

To gain insight into the mechanism of how CDK12 controls epidermal differentiation, we performed Pol II Ser2 ChIP-Seq in control and CDK12 knockdown differentiated cells (Supplementary Table 3A-B). Loss of CDK12 led to a large depletion of Pol II Ser2 at genes where CDK12 bound (Figure 3A). At genes where CDK12 did not bind, there was much less change in Pol II Ser2 binding between control and CDK12 knockdown groups (Figure 3B). There was no change in total levels of Pol II Ser2 in CDK12i cells suggesting that CDK12 specifically phosphorylates Pol II Ser2 on genes that it binds (Figure 3C). These genes include *OVOL1, KLF4, TGM1*, and *GRHL3* where CDK12 loss led to a reduction in Pol II Ser2 binding throughout the gene including the TSS (Figure 3D-G). Since Pol II Ser2 has been correlated with transcription elongation, loss of it may potentially cause total Pol II depletion from the gene body and accumulation at the TSS. To test this, total Pol II ChIP was performed in CTLi and CDK12i differentiated cells. Absence of CDK12 resulted in loss of total Pol II binding from the gene bodies of *OVOL1, KLF4, TGM1*, and *GRHL3* and buildup at the TSS (Figure 4A-B). Importantly no impacts were seen with Pol II binding on house keeping gene *PGK1*, which is a gene that CDK12 does not bind (Figure 4A-B, Supplementary Table 2). This suggests that CDK12 depletion leads to failure to elongate differentiation genes and thus subsequent buildup of Pol II at the TSS of those genes. In addition to promoting elongation through Pol II Ser2 phosphorylation, CDK12 may also be required for elongation factors to bind to differentiation genes. Since we previously demonstrated^37^ that the elongation factor SPT6 was necessary for epidermal differentiation, we compared the SPT6 bound genes to CDK12. Nearly 60% (3,623/6,307) of the genes bound by SPT6^37^ were also bound by CDK12 and ∼38% (1,449/3,855) of the CDK12 gene expression signature overlapped with the SPT6 signature^37^ (Figure 4C-D). These 1,449 overlapped genes were enriched for skin development and keratinocyte differentiation genes suggesting that CDK12 and SPT6 are in the same pathway and that CDK12 may be necessary for SPT6 binding to target genes (Figure 4E). To examine whether CDK12 is necessary for SPT6 binding to differentiation genes, SPT6 ChIP was performed in differentiated CTLi and CDK12i cells. In control cells, SPT6 bound robustly to *OVOL1, GRHL3, KLF4*, and *TGM1* (Figure 4F). In contrast, CDK12 knockdown abolished SPT6 binding to those genes (Figure 4F). The diminished binding is not due to a loss of SPT6 levels as CDK12 knockdown did not alter protein levels of SPT6 (Figure 4G). These results suggest that CDK12 promotes elongation of epidermal differentiation genes through the phosphorylation of Pol II Ser2 and elongation factor binding.

**Figure 3.**
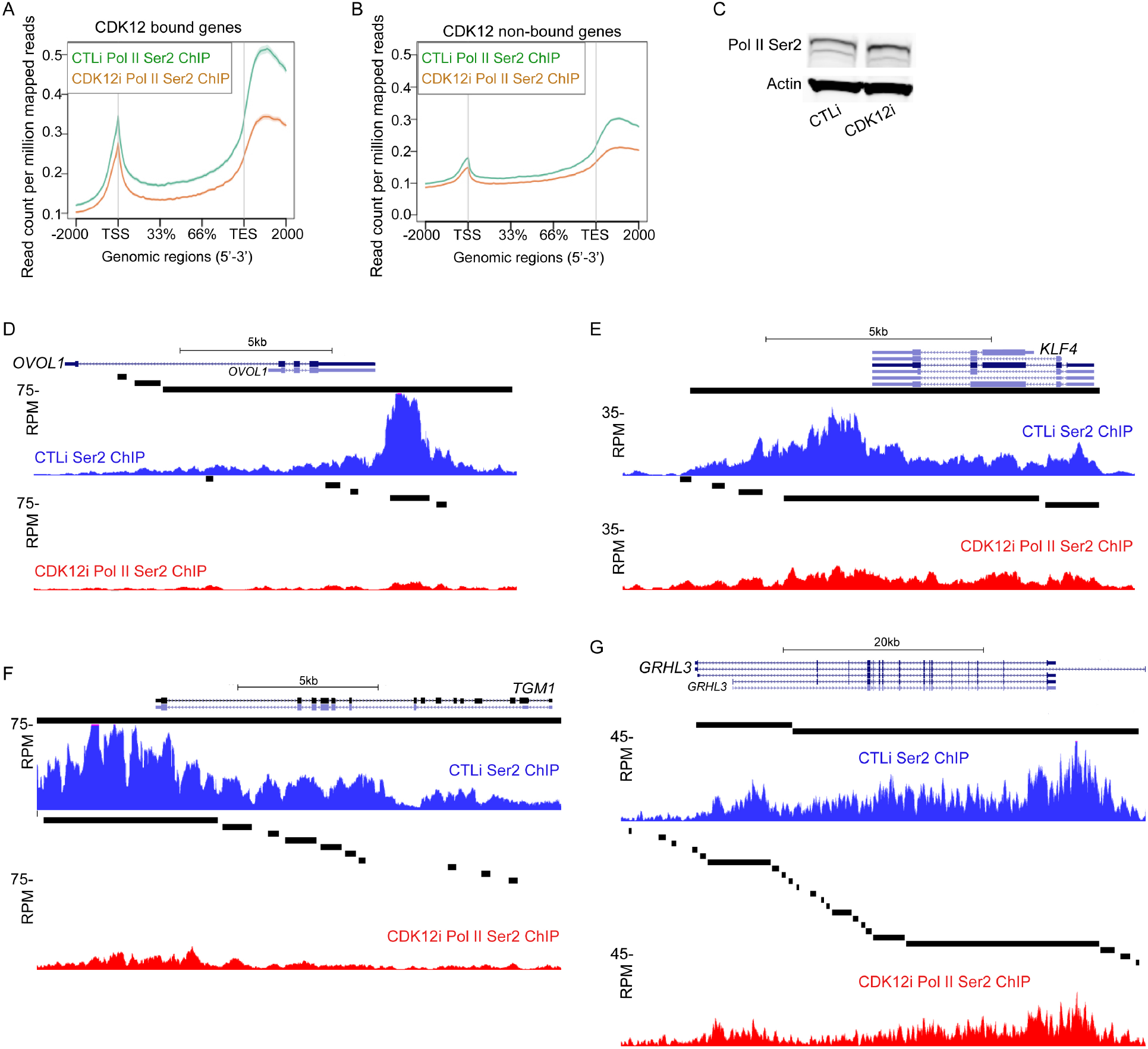
CDK12 is Necessary for Pol II Ser2 Phosphorylation at CDK12 Bound Genes. **(A)** RNA Polymerase II Ser2 (Pol II Ser2) ChIP-Seq at regions bound by CDK12 in CTLi (green) and CDK12i (orange) differentiated cells (Day 3: full confluence and 1.2mM calcium). Y-axis is shown as read count per million reads and X-axis is distance along genes. TSS is transcription start site and TES is transcription end site. Pol II Ser2 ChIP-Seq in CTLi and CDK12i cells were performed in duplicates. **(B)** Metagene plot of Pol II Ser2 ChIP-Seq at regions not bound by CDK12 in CTLi (green) and CDK12i (orange) differentiated cells. **(C)** Western blot of RNA Polymerase II Ser2 (Pol II Ser2) in CTLi and CDK12i differentiated cells. Actin was used as a loading control. N=3. Representative blots are shown. **(D-G)** Gene tracks of *OVOL1* (D), *KLF4* (E), *TGM1* (F), and *GRHL3* (G). Pol II Ser2 ChIP-Seq is shown in CTLi (blue) and CDK12i (red) cells. Y-axis shows reads per million (RPM) and black bar over gene tracks represent significant peaks. X-axis shows position along gene. Source data for Figure 3C can be found in Figure 3-Source Data 1.

**Figure 4.**
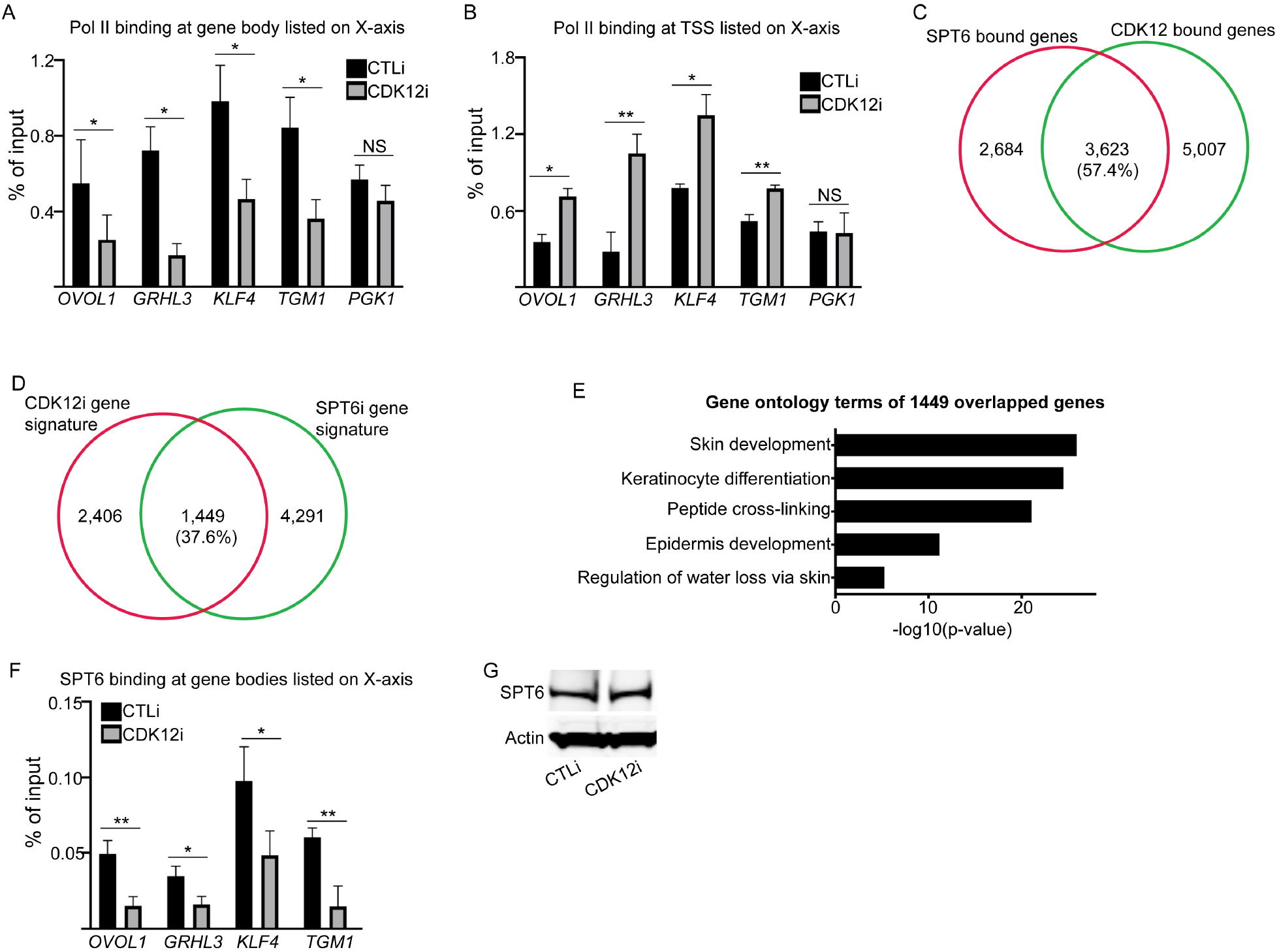
CDK12 is Necessary for SPT6 Binding to Epidermal Differentiation Genes. **(A)** RNA Polymerase II (Pol II) ChIP on CTLi and CDK12i differentiated cells (Day 3: full confluence and 1.2mM calcium). QPCR was used to determine the amount of binding to genes listed on the X-axis. Primers were targeted towards the gene body of each gene. Results are plotted as a percent of input. **(B)** Pol II binding at the transcription start site (TSS) of each gene listed on the X-axis in CTLi and CDK12i differentiated cells. **(C)** Venn diagram of genes bound by SPT6 (SPT6 ChIP-Seq) and CDK12 (CDK12 ChIP-Seq). **(D)** Overlap of the CDK12i gene signature (CDK12i RNA-Seq) with the SPT6i gene signature (SPT6i RNA-Seq). **(E)** Gene ontology of the 1,449 overlapped genes from (D). **(F)** SPT6 ChIP on CTLi and CDK12i differentiated cells (Day 3: full confluence and 1.2mM calcium). QPCR was used to determine the amount of binding to genes listed on the X-axis. Primers were targeted towards the gene body of each gene. Results are plotted as a percent of input. **(G)** Western blot of SPT6 in CTLi and CDK12i differentiated cells. Actin was used as a loading control. Representative blots are shown. N=3 independent experiments for Figure 4A, 4B, 4F,4G. Mean values are shown with error bars=SD. * ***p*** <0.05, ** ***p*** <0.01(t-test). NS=not significant. Source data for Figure 4A, 4B, and 4F can be found in Figure 4-Source Data 1. Source data for Figure 4G can be found in Figure 4-Source Data 2.

## DISCUSSION

The role of CDK12 or CDK13 in adult tissue homeostasis has not been characterized due to the embryonic lethality of murine knockout models^20,21^. This is an important issue that needs to be addressed since CDK12/CDK13 inhibitors have been proposed to be used in combination with PARP inhibitors (to induce synthetic lethality) for breast cancer patients^48,49^. CDK12 knockout leads to embryonic lethality by E6.5^20^. In murine embryonic stem cells CDK12 or CDK13 depletion results in spontaneous differentiation due to loss of pluripotency gene expression^50^. Our results show an opposite phenotype in somatic tissue in that CDK12 but not CDK13 is essential for epidermal differentiation. In the absence of CDK12, regenerated human skin fails to terminally differentiate including a complete absence of stratum corenum formation. CDK12 depletion also led to a thinner epidermis and loss of KI67 positive proliferative cells in the basal layer. This phenotype is likely due to the known role of CDK12 in promoting mammalian proliferation by phosphorylating cyclin E1^51^. Global gene expression profiling of CDK12 knockdown cells showed that 2,725 genes were upregulated and 1,130 genes were decreased in expression. The 1,130 downregulated genes were enriched in keratinocyte differentiation and skin development GO terms. Furthermore, a third of the CDK12 regulated overlapped with the differentiation gene signature. These results indicate that CDK12 controls the differentiation gene expression program.

To gain an understanding of the genes that CDK12 directly regulates to enable differentiation to occur, we mapped the genome-wide binding sites of CDK12. This is also an area of active investigation where there have been differing reports published on the type of target genes that depend on CDK12 for expression. CDK12 has been shown to promote elongation/pocessivity of specific genes such as those involved in the DNA damage response and DNA replication^19,52^. In contrast, Tellier et al. showed that CDK12 controls genome-wide transcriptional elongation while Fan et al. demonstrated that CDK12 was redundant with CDK13 to promote elongation globally^53,54^. These differing results may be due to the differences in immortalized/cancer cell lines used as well as assays used. Our data in primary human skin cells demonstrated that CDK12 bound to 8,630 genes, which is nearly a third of the human genes; however only 490 of those genes were downregulated upon CDK12 loss. Interestingly these 490 CDK12 bound genes that decreased in expression upon CDK12 depletion were enriched for skin barrier and epidermis development genes suggesting that differentiation genes were more sensitive to perturbations in CDK12 levels. This may be due to the cell fate change where differentiation genes are highly induced and thus may be more reliant on transcription elongation. In addition, we had previously shown that many of the differentiation genes including *GRHL3, TGM1*, and *OVOL1* already had paused Pol II at its TSS in epidermal stem and progenitor cells which progressed to active elongation upon induction of differentiation^37^. This suggests that paused genes may be more reliant on CDK12 to promote elongation. Thus during cell fate transitions paused and highly induced genes are more susceptible to perturbations in CDK12 levels.

Next, we wanted to determine how CDK12 impacted Pol II Ser2 levels. On CDK12 bound genes, there was a large reduction in Pol II Ser2 levels in CDK12 knockdown cells whereas there was minimal impact on genes not bound by CDK12. Loss of Pol II Ser2 was found on critical differentiation promoting transcription factors such as *KLF4, GRHL3*, and *OVOL1*. Consistent with other studies^53^, CDK12 depletion led to decreased levels of Pol II Ser2 throughout the gene including the TSS. This in turn led to a reduction of total Pol II found along the gene bodies of *KLF4, GRHL3, TGM1*, and *OVOL1*. The reduction in the gene bodies caused a buildup of total Pol II at the TSS of those genes resulting in a blockade of transcription elongation.

In addition to promoting Pol II Ser2 phosphorylation, CDK12 may also be important for elongation factor binding to target genes. We previously showed that SPT6 is necessary to promote the transcriptional elongation of epidermal differentiation genes and thereby could be dependent on CDK12 for binding to target genes^37^. Supporting this, a majority (57.4%) of the SPT6 bound genes overlapped with CDK12 and nearly 40% of the CDK12 gene expression signature overlapped with SPT6. Most importantly, knockdown of CDK12 resulted in decreased SPT6 binding to its target genes including *KLF4, GRHL3, TGM1*, and *OVOL1*. CDK12 may potentially phosphorylate SPT6 to allow contact with other members of the elongation complex such as Pol II. It was previously shown that THZ531 a small molecule inhibitor of CDK12 and CDK13 impacted SPT6 phosphorylation^55^. Our results here demonstrate that CDK12 and SPT6 act in the same pathway to promote epidermal differentiation whereas other elongation factors are necessary for other aspects of skin physiology. For example, we previously showed that the FACT complex (SSRP1 and SPT16) is not essential for epidermal differentiation but is necessary for transcription elongation of *MAP2K3* to mediate the skin inflammatory response^37,56^. In addition, ELL which is part of the super elongation complex has no impact on differentiation of the skin but is essential for proliferation of epidermal stem and progenitor cells^8,57^. Thus, the elongation factors that have been previously identified through biochemical assays to promote transcription elongation and thought to act in the same pathway actually have different gene targets and functions in tissue. In the future, it will be important to characterize each elongation factor’s function in adult tissue since many are altered in diseases such as cancer^3^.

Our findings suggest that transcription elongation regulation of epidermal differentiation is controlled by CDK12 through two mechanisms. First, CDK12 is required to phosphorylate Pol II Ser2 which allows Pol II to elongate. Second, CDK12 is necessary for the binding of the histone chaperone SPT6 to remodel the chromatin to allow passage of Pol II on DNA. In conclusion, we have found a prominent role for CDK12 in promoting epidermal differentiation through elongation of genes coding for transcription factors and structural proteins.

## Acknowledgements

This work was supported by grants from the National Institutes of Health (NIH R01AR072590, R01AR066530, and R01CA225463) to G.L. Sen.

## METHODS

### Primers and siRNA Sequences

All primers and siRNA sequences can be found in the Primers and siRNA Sequences-Source Data 1.

### Cell Culture

Primary human epidermal keratinocytes (derived from neonatal foreskin) were cultured in EpiLife medium (ThermoFisher:MEPI500CA) supplemented with human keratinocyte growth supplement (ThermoFisher:S1001K) and pen/strep. Proliferating, non-differentiated keratinocytes were cultured in subconfluent conditions. To induce epidermal differentiation, primary human keratinocytes were plated at full confluence in the presence of 1.2mM calcium for 3 days.

### Regenerated Human Skin

For the organotypic skin cultures, one million control, CDK12, or CDK13 knockdown cells were seeded onto devitalized human dermis to regenerate human epidermis^42,44,58,59^. Human dermis was purchased from the New York Firefighters Skin Bank. The seeded cells were raised to the air liquid interface to promote stratification and differentiation. Tissue was harvested 4 days after initial seeding.

### Gene Knockdown

siRNAs targeting human CDK12 (Dharmacon D-004031-01 and D-004031-02) CDK13 (Life Technologies s16398) or control siRNAs (final concentration 10 nM) were transfected into keratinocytes using Lipofectamine RNAiMAX (Thermo:13778-500) reagent according to the manufacturer’s protocol and incubated for 18 hrs.

### RNA Isolation and RT-QPCR

Total RNA from cells was extracted using the GeneJET RNA purification kit (Thermo Scientific: K0732) and quantified using a Nanodrop. One µg of total RNA was reversed transcribed using the Maxima cDNA synthesis kit (Thermo Scientific: K1642). Quantitative PCR was performed using the BioRad LFX96 real-time system. L32 was used as internal control for normalization.

### Western blotting

Twenty microgram of the cell lysates were used for immunoblotting and resolved on 10% SDS-PAGE and transferred to PVDF membranes. Primary antibodies used include beta-actin (Santa Cruz: Sc-47778) at 1:5000, RNA pol II Ser2 (Active Motif: 91115) at 1:1000, and SPT6 (Bethyl: A300-801A) at 1:1000. Secondary antibodies including IRDye 800CW (LI-COR: 926-32212) donkey anti-mouse and IRDye 680RD (LI-COR: 926-68073) donkey anti-rabbit were used at 1:10000.

### Histology and immunofluorescence

Regenerated human skin sections were fixed in 4% paraformaldehyde for 11 min followed by blocking in PBS with 2.5% normal goat serum, 0.3% triton X-100, and 2% bovine serum albumin for 30 min. Primary antibodies used were Loricrin (Abcam: Ab198994) at 1:1000, Filaggrin (Abcam: Ab3137) at 1:200, MKi67 (Abcam: Ab16667) at 1:300, and Keratin 10 (Abcam: Ab9025) at 1:500. The secondary antibodies used were Alexa 555 conjugated goat anti-mouse IgG (Thermo: A11029) or Alexa 488 conjugated donkey anti-rabbit IgG (Thermo: A21206) both at 1:500. Nuclear dye, Hoechst 33342 (Thermo:H3570) was used at 1:1000.

### Hematoxylin and Eosin Staining

Sectioned tissue derived from regenerated human skin was fixed with 10% formalin solution (Sigma HT5012) for 12 minutes. Sections were then dipped in 0.25% Triton-X-100 in PBS for 5 minutes. Haemotoxylin (Vector H-3401) staining was performed for 8 minutes, rinsed in water, and then dipped in acid alcohol (1% HCL in 70% ethanol). After subsequent rinsing, the sections were dipped in 0.2% ammonia water for 1 minute, rinsed again, and then dipped in 95% ethanol. Eosin (Richard-Allan Scientific 71304) staining was performed for 30 seconds followed by 95% ethanol rinsing for 1 minute. Sections were then put into 100% ethanol for 4 minutes followed by 2 minutes in Xylene.

### RNA-sequencing (RNA-seq) and Library Preparation

Control and CDK12i cells were placed in differentiation conditions (full confluence and 1.2mM calcium) for 3 days. Total RNA was isolated using the GeneJET RNA (Thermo: K0732) purification kit and quantified by Nanodrop for control and CDK12i cells. RNA-seq was performed using the Illumina NovaSeq S4 machine at the Institute of Genomic Medicine core facility at UCSD. ∼30 million reads per sample were obtained using pair-ended 100 base long reads.

### Chromatin Immunoprecipitation Sequencing (ChIP-seq), Library Preparation, and ChIP-QPCR

Ten million cells and 5 µg of antibody were used for each antibody pulldown experiment for ChIP^36,43,60^. ChIP was performed using the following antibodies: SPT6 (Bethyl: A300-801A), RNA Pol II (Active Motif: 91151), RNA Pol II Ser2 (Active Motif: 91115), CDK12 (Bethyl: A301-679A and LSBio: LS-A7350), Rabbit IgG (Millipore: 12-370) and mouse IgG (Abcam: Ab18413). Cells for the RNA Pol II/RNA Pol II Ser2 ChIP-QPCR or ChIP-Seq were fixed at a final concentration of 1% formaldehyde. Cells for the SPT6 or CDK12 ChIP-QPCR or ChIP-Seq were fixed in both formaldehyde (1% final concentration, Thermo:28908) and disuccinimidyl glutarate (2 mM final concentration, Thermo: 20593). QPCR results are represented as a percentage of input DNA.

For ChIP-Seq, the ChIP DNA library was prepared using the TruSeq DNA sample prep kit (Illumina). Sequencing was done on the HiSeq 4000 System (Illumina) using single 1 × 75 reads at the Institute for Genomic Medicine Core, UCSD.

### RNA-seq Analysis

Reads were aligned to the GENCODE v19 transcriptome hg19 using TopHat2 with default settings^61^. Differential expression among samples was calculated using ANOVA from the Partek Genomic Suite (Partek Incorporated). Analysis of the read count distribution indicated that a threshold of ten reads per gene generally separated expressed from unexpressed genes, so all genes with fewer than ten reads were excluded from ANOVA analysis. Gene lists for significantly upregulated or downregulated genes were created using FDR < 0.05 and ≥ 2-fold change. Enriched GO terms for RNA-seq differentially expressed gene sets were identified using Enrichr ^62,63^. Heatmaps for the RNA-seq data were generated using Partek’s Genomic Suite (http://www.partek.com/partek-genomics-suite/).

### ChIP-seq Analysis

The ChIP-seq reads were processed by the ENCODE Transcription Factor and Histone ChIP-Seq processing pipeline (https://github.com/ENCODE-DCC/chip-seq-pipeline2) on our local workstation. The reads were first trimmed based on quality score before alignment to reference hg19; Upon alignment and deduplication, the peak-calling was then carried out by MACS2.2.4 with a cutoff q-value of 0.05^64,65^. The heatmaps for the ChIP-Seq data were generated using ngs.plot^66^. Gene tracks were visualized using UCSC genome browser along with annotation tracks.

## DATA AVAILABILITY

The datasets generated from this study including RNA-Seq and ChIP-Seq data has been deposited in GEO (GSE166407) with reviewer token ktstacckfjuhxsj.

---

**Supplementary Figure 1.**
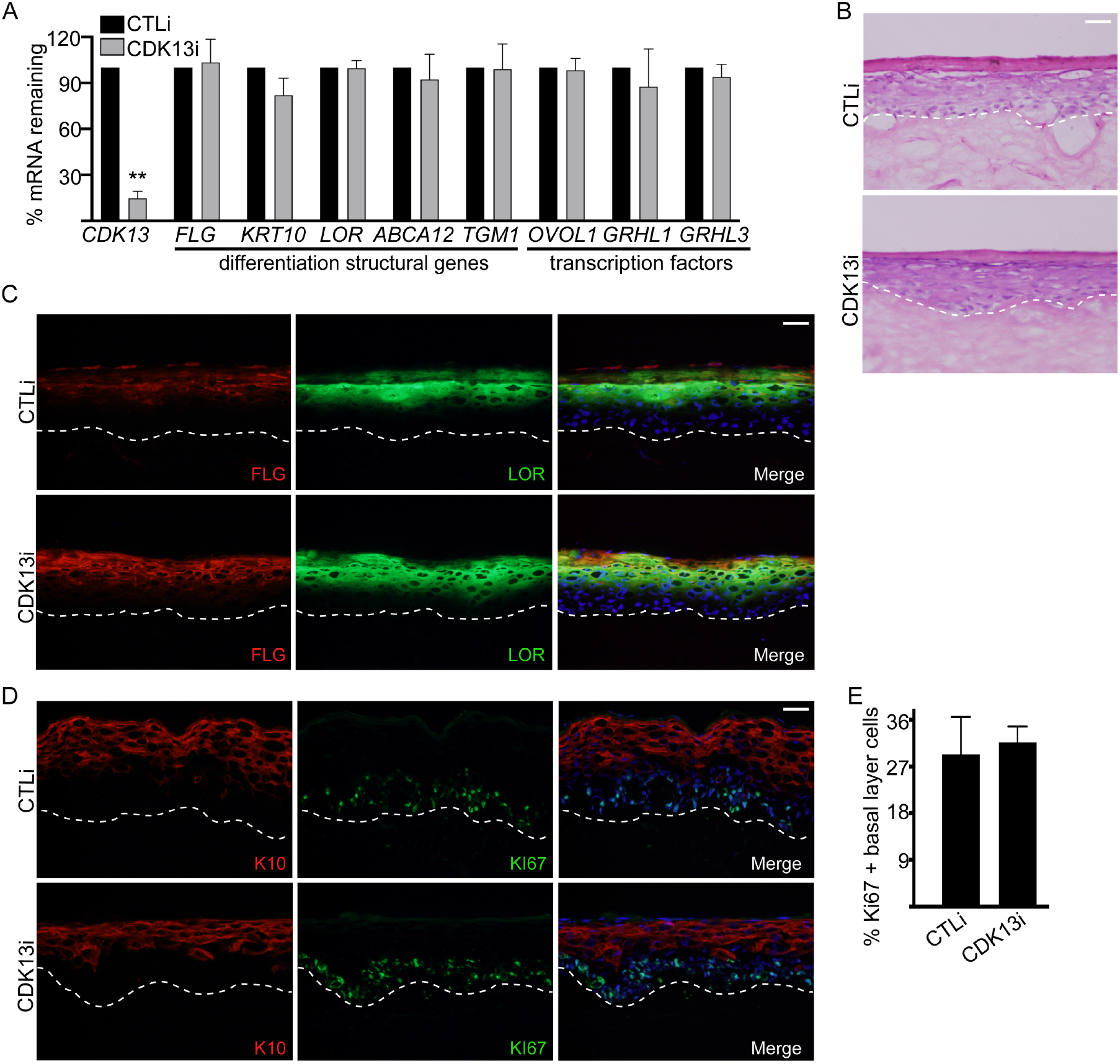
CDK13 is Not Essential for Epidermal Growth or Differentiation. **(A)**Primary human keratinocytes were knocked down with control (CTLi) or CDK13 (CDK13i) siRNAs and placed in differentiation conditions for 3 days (full confluence and 1.2mM calcium). The mRNA levels of each gene (X-axis) were measured using RT-qPCR. QPCR results were normalized to *L32* levels. **(B)** Regenerated human skin using three-dimensional organotypic cultures made from CTLi or CDK13i cells were harvested after 4 days of culture. Hematoxylin and eosin staining of regenerated CTLi and CDK13i human skin. **(C)** Immunostaining of late differentiation proteins loricrin (LOR: green) and filaggrin (FLG: red) are shown (Day 4 of organotypic culture). Nuclei are shown in blue (Hoechst staining). **(D)** Immunostaining of CTLi or CDK13i regenerated human skin (Day 4) with early differentiation marker keratin 10 (K10: red) and proliferation marker KI67 (green). **(E)** Quantification of the percentage of Ki67 positive cells in the basal layer from (D). The dashed white lines denote the basement membrane zone and scale bar=20μm for (B-D). N=3 independent experiments for Supplementary Figure 1. Mean values are shown with error bars=SD. ** ***p*** <0.01 (t-test). Source data for Supplementary Figure 1A, 1E can be found in Supplementary Figure 1-Source Data 1.

**Supplementary Figure 2.**
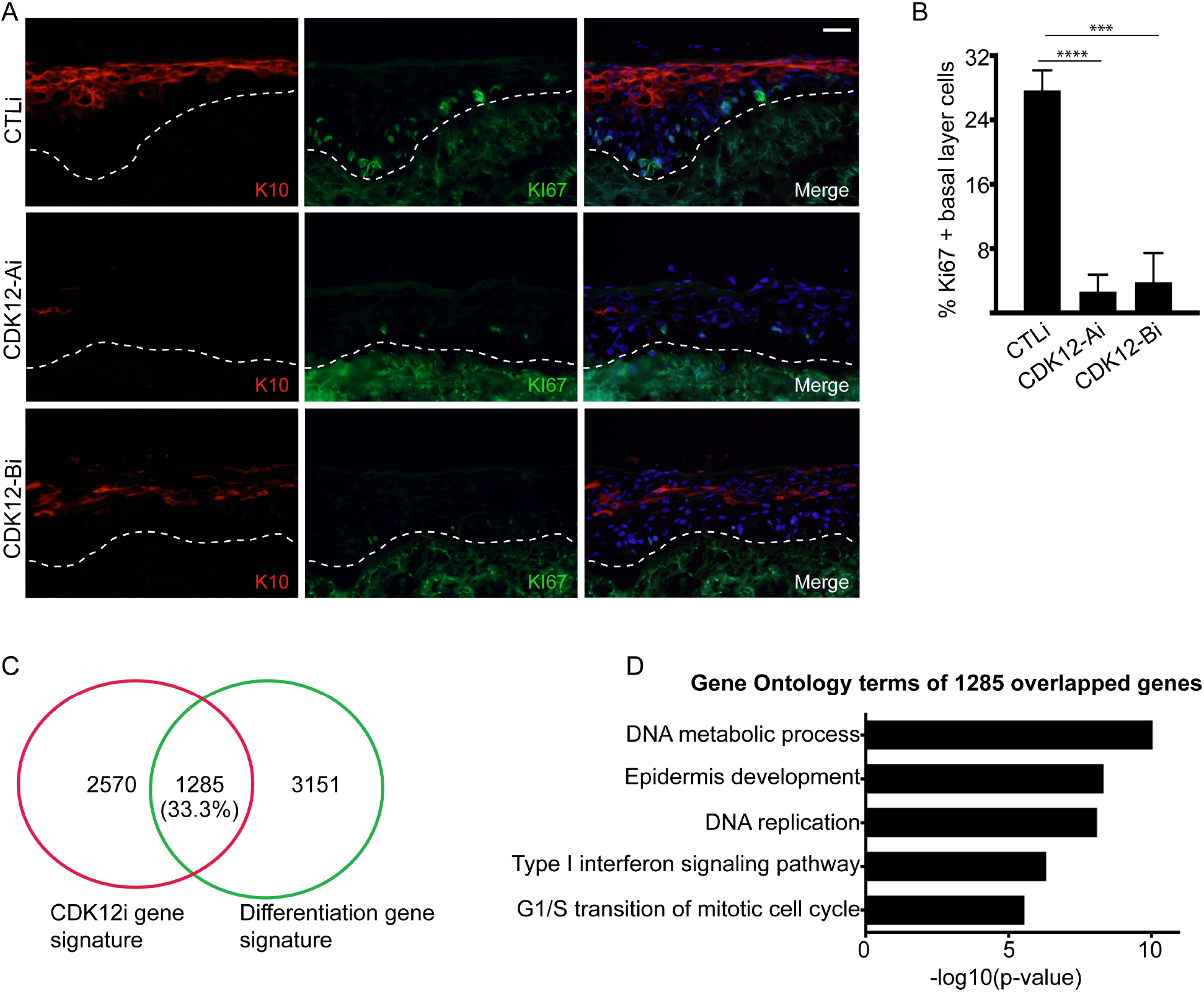
CDK12 is Necessary for Epidermal Growth and Differentiation. **(A)** Immunostaining of CTLi or CDK12i regenerated human skin (Day 4) with early differentiation marker keratin 10 (K10: red) and proliferation marker KI67 (green). The dashed white lines denote the basement membrane zone and scale bar=20μm. N=3. **(B)** Quantification of the percentage of Ki67 positive cells in the basal layer from (A). N=3. **(C)** Venn diagram of the overlap between the CDK12i gene expression signature (CDK12i RNA-Seq) with the differentiation gene signature (genes differentially expressed between undifferentiated and differentiated primary human keratinocytes). **(C)** Gene ontology terms of the 1,285 overlapped genes from (C). Source data for Supplementary Figure 2B can be found in Supplementary Figure 2-Source Data 1.

**Supplementary Figure 3.**
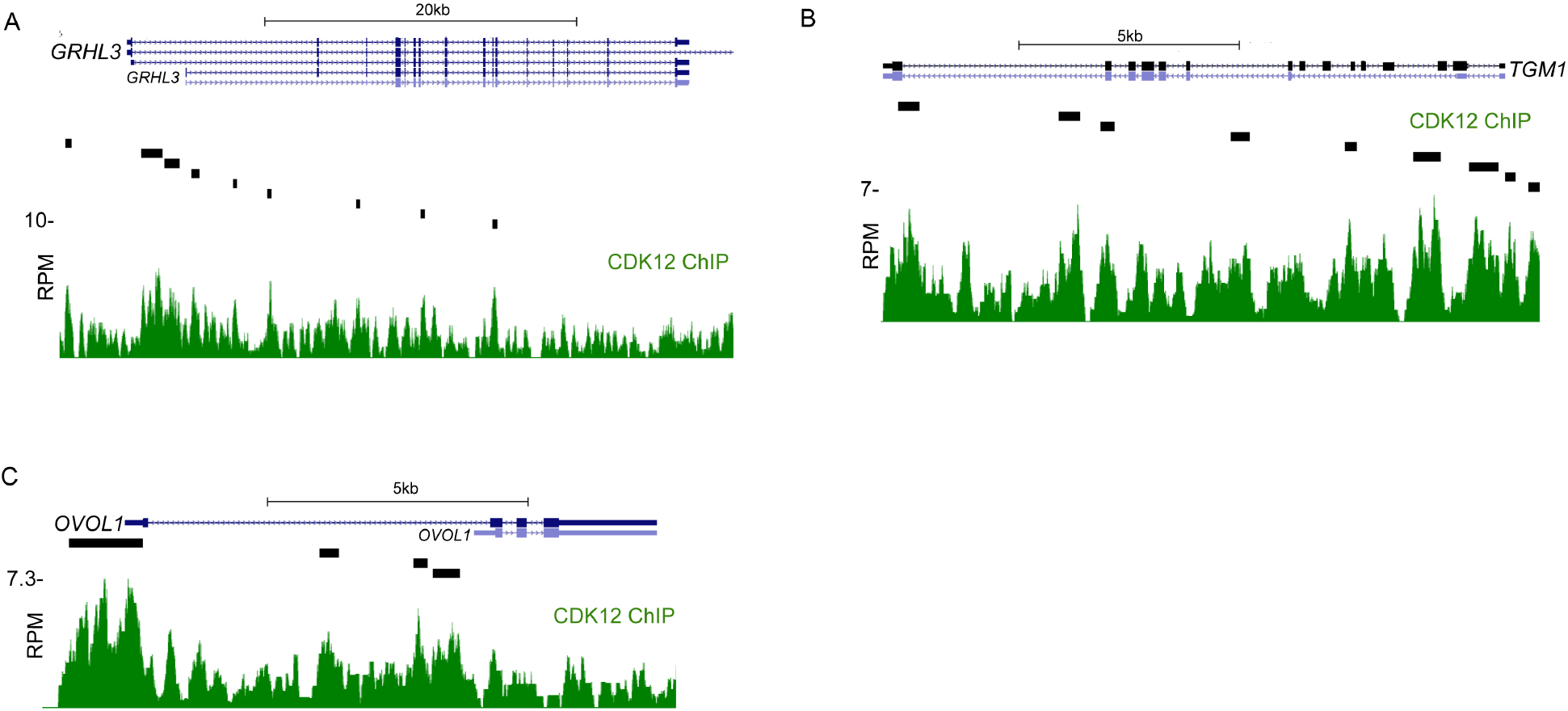
CDK12 Binds to Genes that Code for Epidermal Differentiation Transcription Factors and Structural Proteins. **(A-C)** Gene tracks of *GRHL3* (A), *TGM1* (B), and *OVOL1* (C). CDK12 ChIP-Seq is shown in green. Y-axis shows reads per million (RPM) and black bar over gene tracks represent significant peaks. X-axis shows position along gene.

